# Hexose transporters CsHT3/16 involved in post-phloem transport and affected cucumber fruit development

**DOI:** 10.1101/2024.01.25.577232

**Authors:** Jintao Cheng, Suying Wen, Kexin Li, Yixuan Zhou, Mengtian Zhu, H. Ekkehard Neuhaus, Zhilong Bie

**Author notes:** To whom correspondence should be addressed Jintao Cheng, Zhilong Bie. National Key Laboratory for Germplasm Innovation and Utilization for Fruit and Vegetable Horticultural Crops, College of Horticulture and Forestry Sciences, Huazhong Agricultural University, Wuhan 430070, Hubei Province, China. these authors contributed equally to this paper (Jintao Cheng, Suying Wen).

## Abstract

Hexoses are crucial for plant growth and fruit development. However, the role of hexose transporters in post-phloem sugar transport and maintenance of cellular sugar homeostasis in rapidly growing fruits, such as cucumber, is not yet fully understood. To clarify the impact of hexose transporters in cucumber fruits, we conducted systematic analyses of their tissue expression, localization, transport characteristics and physiological functions. The study revealed that *Cs*HT3, *Cs*HT12 and *Cs*HT16 are the primary hexose transporter genes expressed in cucumber fruit. During the ovary and young fruit stages, *Cs*HT3 and *Cs*HT16 were located in the SE/CC system, but as the cucumber fruit developed and expanded, both transporters shifted to phloem parenchyma cells. The knock-out mutants of *CsHT16* display shorter fruits with a larger circumference, likely due to impaired homeostasis of sugars and hormones. Simultaneously reducing the expression of *CsHT3*, *CsHT12* and *CsHT16* leads to decreased fruit size. Conversely, overexpression of *CsHT3* results in increased fruit size and higher fruit sugar levels. Our data suggest that *Cs*HT16 plays an important role in maintaining sugar homeostasis to shape the fruit, while *Cs*HT3, *Cs*HT12 and *Cs*HT16 together determine the carbohydrates requirement of the enlarged cucumber fruit.

## Introduction

The past two decades have seen extensive research on source-sink relationships in plants aiming to enhance crop productivity and quality. This, in turn, relies on effectively regulating the metabolic interplay between leaves and storage organs (Fernie *et al*., 2020). Cucumber fruits exhibit rapid growth, as their size can increase by over 10 times in just a few days. These rapidly growing fruits impact leaf growth at nodes nine and ten, and up to nine source leaves simultaneously transport diverse forms of carbohydrates to the cucumber fruit (Ells J.E, 1983; Pharr *et al*. 1985). Latter findings indicate that developing cucumber fruits act as a significant sink for carbohydrates derived from source leaves (Ho, 1988; Marcelis, 1996).

The molecular mechanism responsible for maintaining sink strength in cucurbitaceous fruits remains largely unknown. Nevertheless, various studies suggest that sugar unloading, transfer, and storage within fruits may play a pivotal role in determining sink strength ( Bihmidine *et al*., 2013; Bermudez *et al*., 2014). Generally, loading of carbohydrates into the phloem occurs either via the symplastic or apoplastic pathway, or at both routes at the same time (Lalonde *et al*., 2003; Aoki *et al*., 2006). For symplastic loading, carbohydrates must diffuse through the plasmodesmata from the sieve element/companion cell (SE/CC) system to the surrounding parenchyma cells. In contrast, apoplastic loading relies on the uptake of sugars via SUC2/SUT2 type sugar importers, which is driven by the proton gradient across the plasma membrane ( Patrick, 1997; Hedrich *et al*., 2015). In sinks that typically accumulate sugars at high concentrations, phloem unloading occurs via the apoplastic pathway, which allow to maintain the static pressure gradient between the SE/CC system and the surrounding parenchyma cells (Lalonde *et al*., 2003; McCaskill & Turgeon, 2007). Interestingly, although the cucumber fruit do not show accumulation of sugars at high concentrations, the corresponding sugar export from the phloem occurs via the apoplastic pathway (Hu *et al*., 2011).

Typically, cucumber, as other Cucurbitaceous, are raffinose family oligosaccharide (RFO)-transporting plants that translocate, besides low levels of sucrose, stachyose and raffinose as major types of sugars in the phloem (Mitchell *et al*., 1992a,b). Although RFOs are the main assimilate form in the phloem, RFOs in cucumber fruits are only present at low levels, while sucrose and hexoses represent major sugars in the peduncles and fruit tissues. Latter finding leads to the conclusion that sucrose and hexose, rather than stachyose or raffinose, are translocated into the cucumber fruit (Gross & Pharr, 1982; Handley *et al*., 1983; Hu *et al*., 2009; Pharr *et al*., 1977), making the hydrolyzation of RFOs into sucrose and galactose by α-galactosidase mandatory prior to sugar unloading into the fruit (Dai *et al*., 2006). Sucrose released from latter process is possibly directly transported, or converted to glucose and fructose by cell-wall invertase prior to import into parenchymal cells (Ruan, 2014). The second product of RFOs hydrolyzation is galactose, which is converted to glucose ( Dai *et al*., 2006, 2011).

Different types of sugar transporters are usually involved in phloem unloading which occurs in storage organs. SWEET proteins are most likely responsible for unloading sucrose from the phloem in sink organs (Chen *et al*., 2012; Frank *et al*., 2012; Jian *et al*., 2016; Mizuno *et al*., 2016). For post-phloem transport sucrose transporters of the SUT/SUC clade (Chincinska *et al*., 2008; Milne *et al*., 2017; Peron *et al*., 2017) and MST type hexose transporters (McCurdy *et al*., 2010; Cheng *et al*., 2015a; Cheng *et al*., 2015b) are required, as all of these act as proton-driven importers (Pommerrenig *et al*., 2018). Accordingly, in sinks, such as fruits from tomato, which mainly accumulate glucose and fructose, hexose transporters have been identified as key players for sugar accumulation ( Afoufa-Bastien *et al*., 2010; McCurdy *et al*., 2010).

In cucumber fruits the concentrations of the monosaccharides glucose and fructose are considerably higher than that of the disaccharide sucrose (Handley *et al*., 1983; Hu *et al*., 2009). This observation is in line with the markedly high activity of the cell-wall invertase in cucumber fruits (Hu *et al*., 2011). Therefore, most of the sucrose released from the transport phloem into the fruit apoplastic space is possibly converted into glucose and fructose prior to transport into parenchymal and flesh cells. Hence, hexose transporters play an important role in this step.

Because cucumber fruits represent strong sinks, exhibit the apoplastic sugar unloading and show a high cell-wall invertase the activity of hexose transporters may play an important role in providing carbon moieties and energy for cellular metabolism. As MST type hexose transporters are pumping their substrates against a concentration gradient they may also be involved in the regulation of the osmotic potential gradient between source and sink which is required to drive mass transport carbohydrates to the fruit.

However, the functional mechanism of such hexose transporters (in particular their substrate specificities), and their exact role in sugar transport and cucumber fruit development are still obscure. To raise answers to these questions we thoroughly analyzed the expression patterns of the hexose transporter genes (*HT*) of the *SPT*/*HT* subfamily during cucumber fruit development. These analyses allowed to identify three *HTs* (namely *HT3*, *HT12* and *HT16*) as highly expressed in cucumber fruits. By combining bioinformatics analyzes, qRT-PCR, subcellular localization, yeast heterologous expression system, immunohistochemical localization and reverse genetics the functions of these genes/proteins in cucumber fruit development were deciphered. In summary, our findings indicate that these proteins are hexose transporters and locate to the plasma membrane of phloem-specific cells. Altered expressions of *HTs* affect the fruit shape, fruit size, and sugar content in cucumber.

## Results

### Identification and phylogenetic analysis of *Cs*HTs in cucumber

To identify the cucumber *HT*/*STP* genes, we blasted 14 *Arabidopsis* STP (*At*STP) proteins against the predicted proteome of the cucumber genome sequence. A total of 16 HTs were identified in *Cucumis sativus* (*Cs*HTs) on the basis of their similarities to *At*STPs (Supplemental Fig. S1, Supplemental Table S2). The *CsHTs* were distributed on 6 out of 7 chromosomes based on the genomic sequences of cucumber (ssp. Chinese long). Most genes are distributed on the third and sixth chromosome, which harbor five and three genes, respectively. Two genes were present on each of the chromosomes 1, 2, 4, and 5 and no *CsHT* was found on chromosome 7 (Supplemental Fig. S2, Supplemental Table S2). Four genes (*CsHT6*, *CsHT7*, *CsHT8*, and *CsHT10*) and two pairs of genes (*CsHT3*/*CsHT13* and *CsHT2*/*CsHT16*) were tightly linked on the respective chromosome (Supplemental Fig. S2). The open reading frames (ORFs) of the *CsHTs* ranged from 1,500 bp to 1,620 bp, encoding polypeptides that are between 500 aa to 540 aa long (Supplemental Table S2). The genomic structures of the 16 *CsHTs* indicate that their intron and exon numbers vary from one intron/two exons (*CsHT11*) to four introns/five exons (*CsHT7* and *CsHT9*) (Supplemental Table S2).

We constructed a phylogenetic tree to visualize the evolutionary relationships between *Cs*HTs and *At*STPs (Supplemental Fig. S1). The results indicated that *Cs*HT1 and *Cs*HT11 are most closely related to each other and homologous to *At*STP9, *At*ST10, and *At*STP11. *Cs*HT12 and *Cs*HT16 are highly homologous to *At*STP1 and *At*STP12. *Cs*HT3, *Cs*HT4, and *Cs*HT15 are homologous to AtSTP5, AtSTP14, and AtSTP7, respectively. Lastly, *Cs*HT5–*Cs*HT10 are all highly homologous to *At*STP13 (Supplemental Fig. S1).

### Spatiotemporal expression analysis of *CsHTs*

To search for the *CsHTs* genes that are expressed in cucumber fruits, we performed semi-RT-PCR to detect the transcripts of *CsHTs* in different organs of cucumber. Out of in total 16 *HT*s, 6 (namely *CsHT3*, *CsHT4*, *CsHT9*, *CsHT12*, *CsHT15*, and *CsHT16*) were found in cucumber fruits (Fig. 1A and B). To identify the relative expressions of these genes in fruits, we performed qRT-PCR and discovered that the relative transcript levels of *CsHT3*, *CsHT12*, and *CsHT16* are considerably higher than those of other *HT* genes, and that they exhibited similar expression patterns during fruit and ovary development (Figs. 1B and C). In fruits/ovary, *CsHT3*, *CsHT12*, and *CsHT16* maintained high relative transcript levels during the late ovary developmental stages (S9–S12) and fruit enlargement stages (3–9 day after anthesis (DAA)), with a low point at 0 DAA (Fig. 1C).

**Fig. 1.**
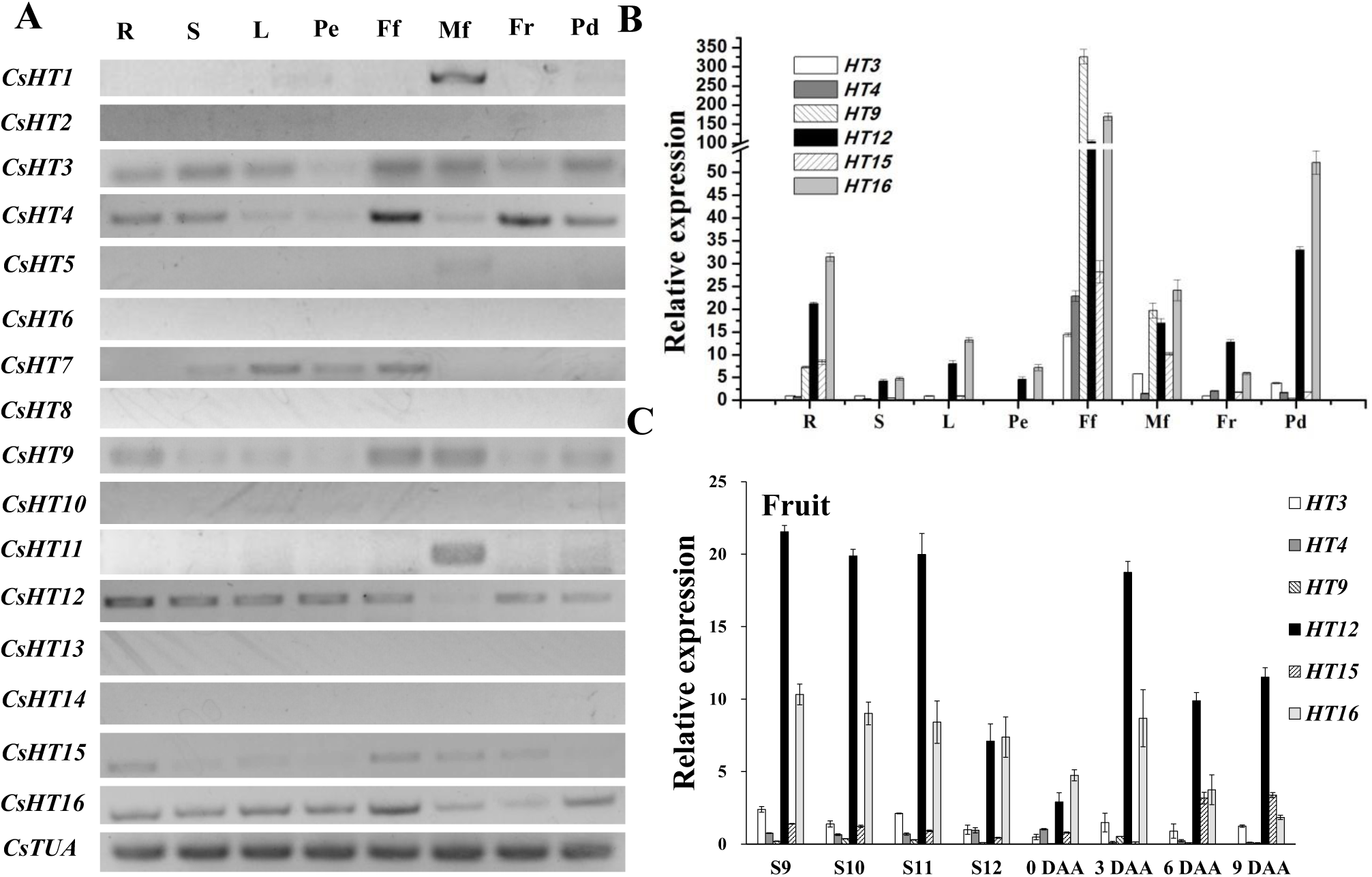
Spatiotemporal expression analysis of *CsHTs*. (A) semi-RT-PCR detection on the expressions of 16 *CsHTs* in different organs of cucumber, *CsHTs* run in 35 cycles, TUA run in 25 cycles; (B)-(C) qRT-PCR detection on the expressions of six *CsHTs* in different cucumber organs (B) or different developmental stages of fruit (C). R, root; L, stem; Pe, petiole; Ff, female flower; Mf, male flower; Fr, fruit; Pd, peduncle; Stages 9–12, four developmental stages of ovary; DAA, day after anthesis; TUA was used as the control. Error bars represent the SE for four technical replicates of three biological replicates.

### Functional characterization analysis of *Cs*HT9, 12, 15, and 16 in yeast

The transport properties of CsHT3 and CsHT4 are already analyzed in the bakeŕs yeast system (Cheng *et al*., 2015a). Thus, we exploited this recombinant system for functional analyses of *Cs*HT9, *Cs*HT12, *Cs*HT15, and *Cs*HT16, which was carried out in comparison to sugar transport into the bakeŕs yeast mutant EBY.VW4000, lacking all endogenous sugar transport proteins.

Growth-based complementation and substrate specificity assays indicated that *Cs*HT9, *Cs*HT12, and *Cs*T16 can restore bakeŕs yeast growth in glucose, fructose, galactose, and mannose. In contrast, *Cs*HT15 cannot restore yeast growth in glucose and galactose, but partly restored the growth in fructose and mannose (Fig. 2A). The time course in media with glucose as sole carbon source showed that the growth rates of yeast which were transformed with either *Cs*HT9, *Cs*HT12, or *Cs*HT16, were linear and markedly higher than those cells carrying the empty vector control (Fig. 2B).

**Fig. 2.**
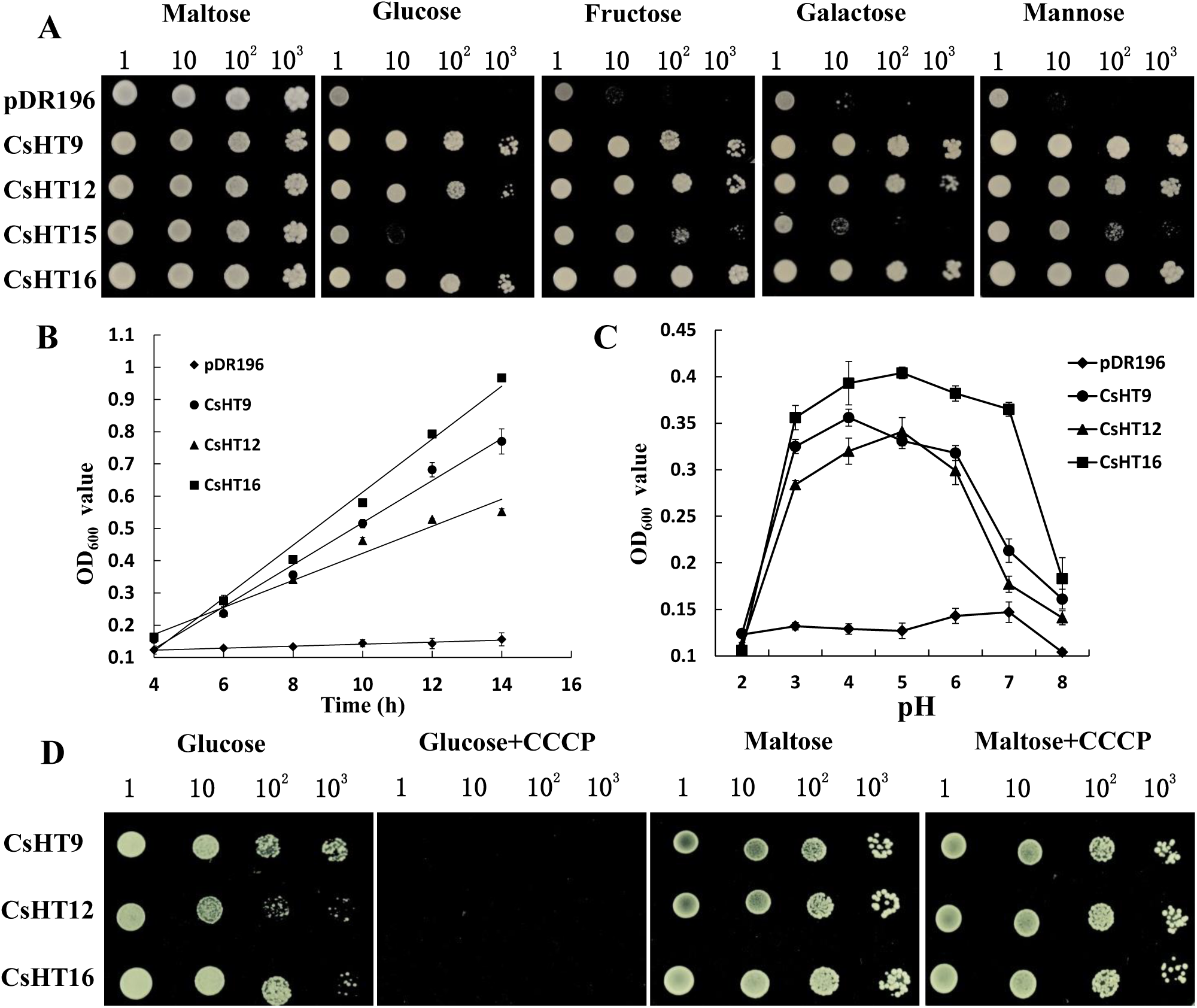
Functional characterization analysis of *Cs*HTs in yeast. (A) Growth complementation on *different* monosaccharides of the hexose transporter-deficient yeast by *CsHT* expression. Four independent transformant lines of EBY.VW4000 expressing *CsHT9*, *CsHT12*, *CsHT15*, and *CsHT16* were grown on maltose for 2 days or on different monosaccharides as the sole carbon source for 4 days. The transformant line with the empty vector pDR196 was used as control. (B) Time course of yeast growth (OD_600_ value) *media with glucose as sole carbon source.* (C) *pH dependence of yeast growth* (OD_600_ value) *in media with glucose as sole carbon source.* (D) Effect of metabolic inhibitor (CCCP) on yeast growth*. Values in (B) and (C) are mean ± SE (n = 3)*.

The transformed bakeŕs yeast cells exhibited a pH-dependent growth, with an optimum range of 3.0 to 6.0 (Fig. 2C). The presence of proton gradient uncoupler carbonyl cyanide *m*-chlorophenylhydrazone (CCCP) provoked a severely inhibited growth of yeast cells expressing *Cs*HT9, *Cs*HT12, and *Cs*HT16 in glucose medium (Fig. 2D). A control experiment showed that yeast cells grew well on the maltose medium containing CCCP. In sum, *Cs*HT9, *Cs*HT12, and *Cs*HT16 exhibit glucose, fructose, galactose, and mannose transport activities, that this transport is energy dependent and that it occurs in symport with protons.

### Subcellular localization of *Cs*HT9, 12, and 16

Our previous work showed that both, *Cs*HT3 and *Cs*HT4 locate to the plasma membrane (Cheng *et al*., 2015a). To determine now the subcellular localization of *Cs*HT9, 12, and 16, that are functional (Fig. 2) and expressed in fruits (Fig. 1), we generated fluorescing reporter proteins coding for *CsHT9-GFP*, *CsHT12-GFP*, and *CsHT16-GFP*, under control of the cauliflower mosaic virus 35S promoter. Each construct was co-expressed with the mCherry-labeled plasma membrane marker (PM-rk; *Arabidopsis* information resource stock, No. CD3-1007) in *Nicotania benthamiana* leaves by infiltration with Agrobacteria.

Confocal analyses of *Agrobacterium*-infiltrated *N. benthamiana* leaves and of protoplasts isolated from infiltrated leaves demonstrated that *Cs*HT9, *Cs*HT12, and *Cs*HT16 locate to the plasma membrane (PM) (Fig. 3A and Fig. S3). Furthermore, plasma membrane localization of *Cs*HT16-GFP has been confirmed on cucumber protoplasts after transient expression of the corresponding DNA construct. Therefore, the data above indicate that three transporters, *Cs*HT9, *Cs*HT12, and *Cs*HT16 located to the plasma membrane.

**Fig. 3.**
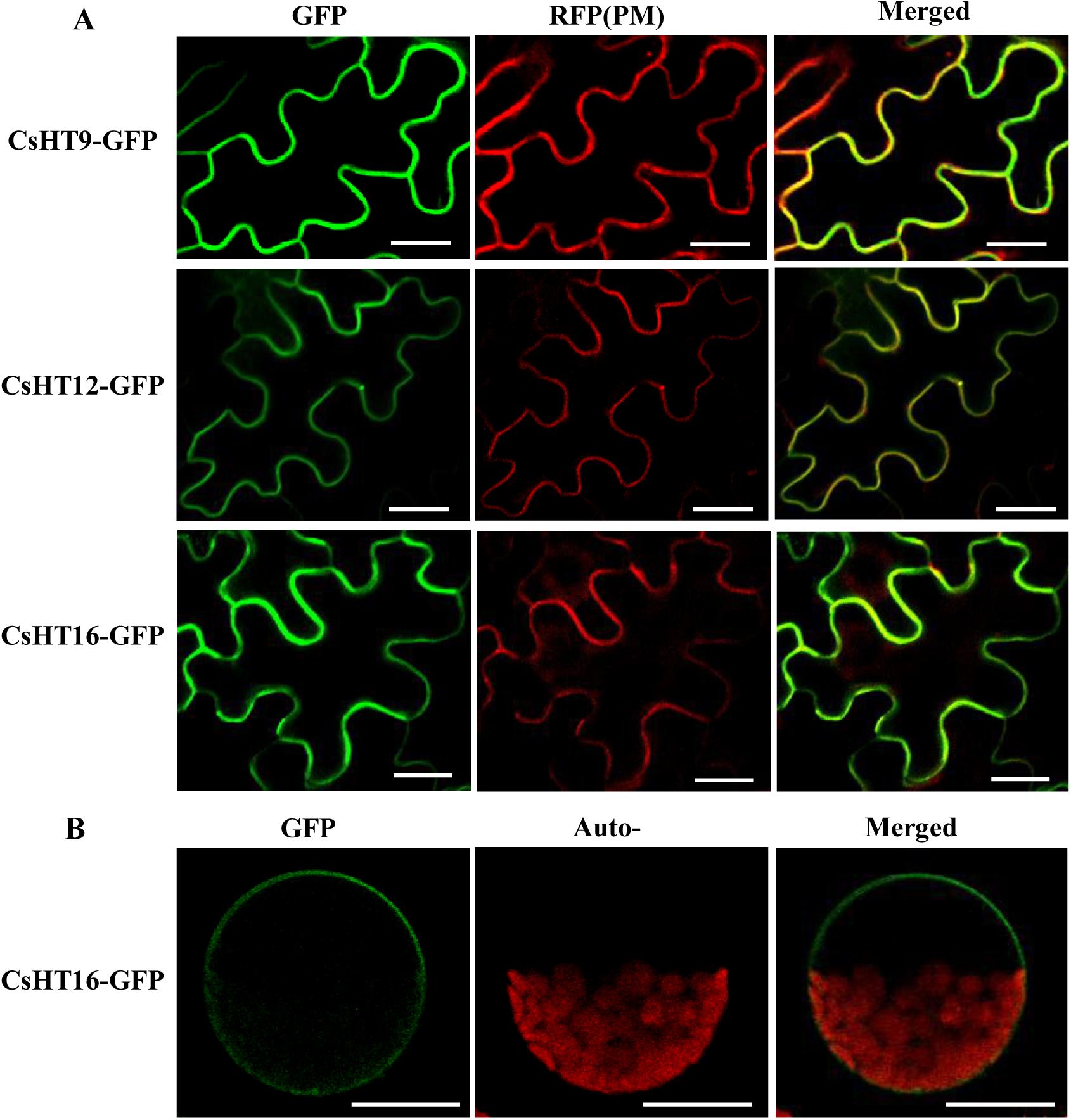
Subcellular localization of *Cs*HT-GFP. (A) Co-localization of *Cs*HT9-GFP, *Cs*HT12-GFP, or *Cs*HT16-GFP with PM-mCherry in tobacco epidermal cells. (B) *Cs*HT16-GFP located in the plasma membrane of cucumber protoplast. The laser scanning confocal microscopy images show fluorescence and merged images. The chlorophyll autofluorescence (Auto-) in (B) is also presented. GFP, green fluorescence protein; RFP, red fluorescence protein; Bar = 20 µm.

### *Cs*HT3 and *Cs*HT16 expressions shift from the external phloem to internal phloem of the bicollateral vascular bundle during ovary-to-fruit transition

To determine the spatiotemporal expression/location pattern of *Cs*HT3 and *Cs*HT16 during ovary-to-fruit transition, immunohistochemistry analysis was performed. To test the specificity of the antisera used Western blot analyses were carried out confirming that the anti-CsHT3 antiserum and anti-CsHT16 antiserum were selectively bound to *Cs*HT3 or *Cs*HT16, respectively (Fig. 4A).

**Fig. 4.**
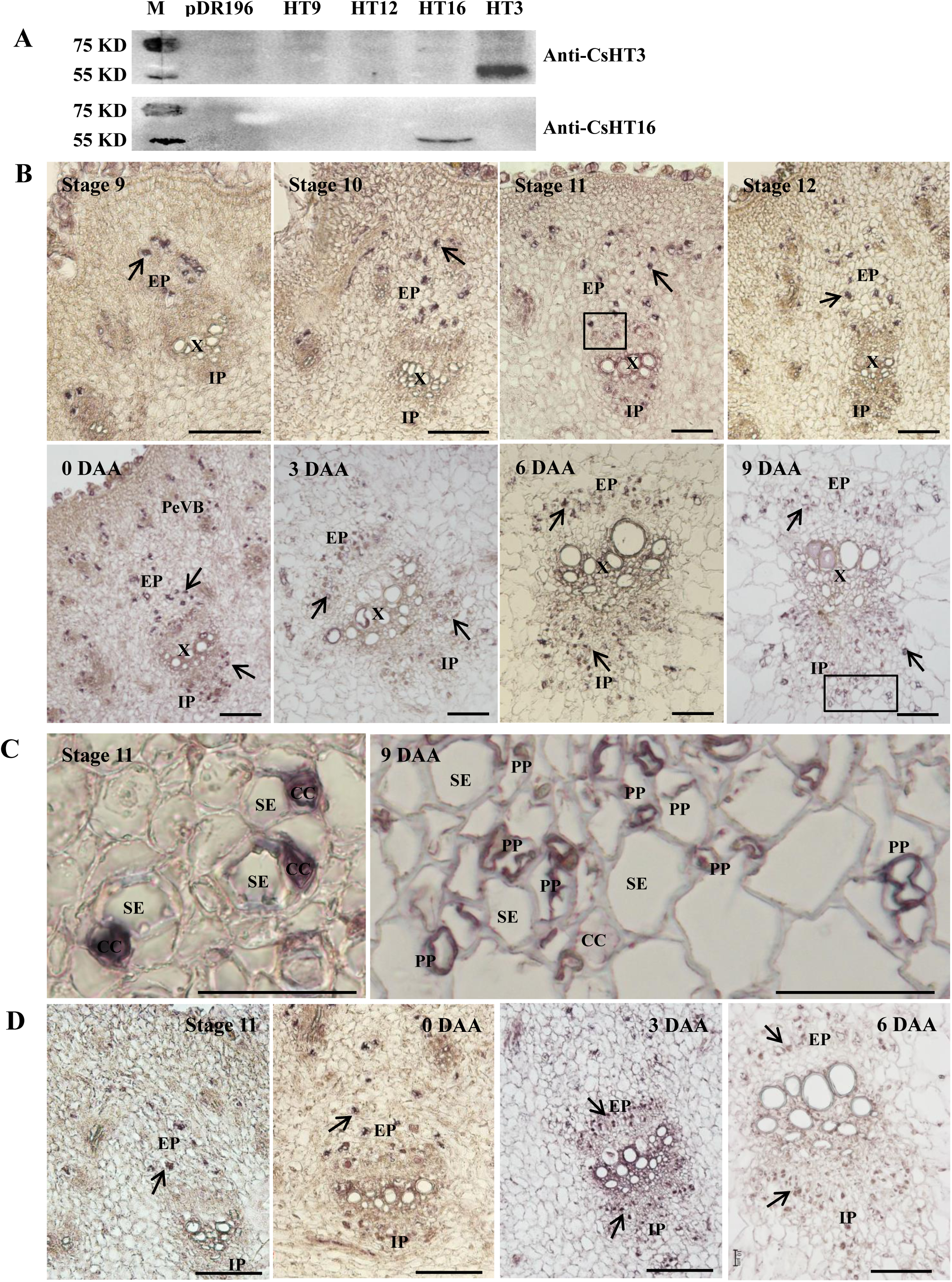
Immunolocalization of *Cs*HT3 (B and C) and *Cs*HT16 (D) in cucumber fruit at different development stages. (A) Western blot to test the quality and specificity of anti-*Cs*HT3 antiserum or anti-*Cs*HT16 antiserum. A specific band was detected in cells expressing *CsHT3 or CsHT16* when anti-*Cs*HT3 or anti-*Cs*HT16 was used, respectively. (B) and (D) Cross-sections of fruits. Bar = 100 µm. (C) Close-ups of the boxes in (B). Bars = 50 µm. PF, phloem fiber; EP, external phloem; IP, internal phloem; X, xylem; SE, sieve element; CC, companion cell; PeVB, periphery vascular bundle; PP, phloem parenchyma cell; Arrows, phloem-specific cells.

For the immunohistochemical localization, histological sections from different developmental stages of ovary and fruits were used. The results showed that *Cs*HT3 and *Cs*HT16 specifically located to the external phloem cells of ovaries (Figs. 4B and C) and after flowering, *Cs*HT3 and *Cs*HT16 appeared on both, external and internal phloem cells (Figs. 4B and C). A partially enlarged image of the ovary phloem cells (S11) and fruits (9 DAA) revealed that *Cs*HT3 was specifically localized in CCs of the phloem system in the ovary (Fig. 4C). However, at later stages *Cs*HT3 was present in the fruit phloem parenchyma cells (Fig. 4C). No signals can be detected in the histological sections incubated with their respective pre-immune sera (Figs. S4).

### Knocking out of the *CsHT16* gene caused shorter and bigger fruits due to impacting hormone signaling

In order to study the physiological function of *Cs*HT12 and of *Cs*HT16, which showed highest transcript levels in fruits (Fig. 1), we constructed gene editing vectors of *CsHT16* and *CsHT12* using CRISPR/Cas9 technology followed by Agrobacterium mediated transformation into the wild-type cucumber variety. Although several attempts for gene silencing were made only one gene edited cucumber line of *Cs*HT16 was identified and for *Cs*HT12 we failed to create a corresponding gene edited mutant.

The insertion of a T base in the second exon caused a stop codon error of *CsHT16*, resulting in a functional deficiency (Fig. 5A). After multiple generations of self-breeding and genotyping a homozygous trans-free *ht16* gene edited cucumber line was identified. Through phenotype observation and field testing of a large number of plants, it became apparent that *ht16* mutants exhibited shorter, but bigger (in diameter) fruits when compared to control plants (Fig. 5B). The length of both, fruits and carpopodia of *ht16* mutants at 20 DAA were significantly lower than those of wild types, while the fruit diameter of *ht16* mutant was significantly larger than that of the WT (Fig. 5C).

**Fig. 5.**
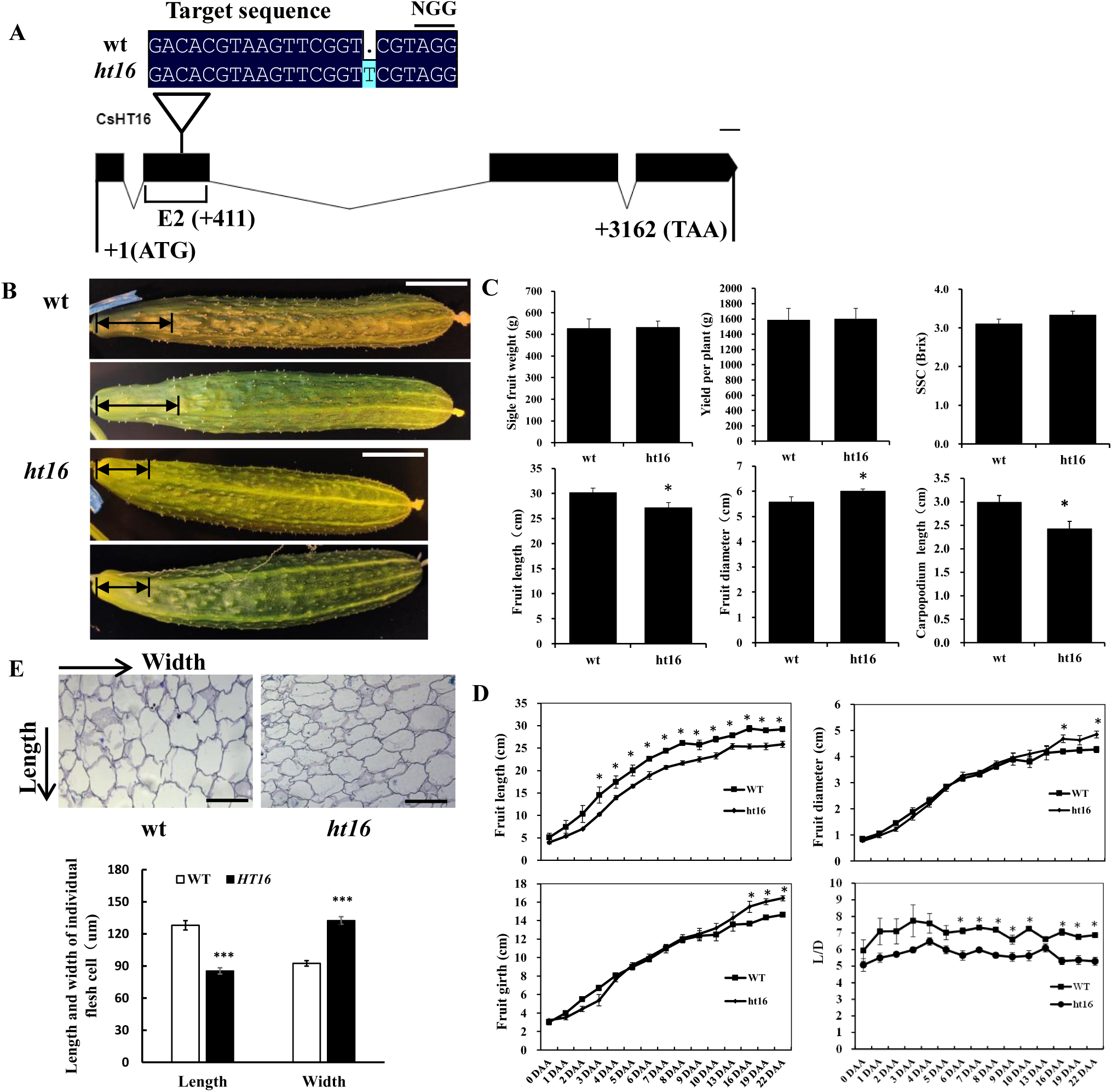
Phenotype of *CsHT16* knock out lines. (A) Exon-intron structure of the *Cs*HT16 gene and gene editing type. Black boxes indicate exons, black lines between boxes indicate intron sequences. Triangles mark gRNA location and gene editing type. Numbers below exons indicated base pair positions of gRNA and start and stope coding position. (B) Fruit photos (16 DAA) of wt and *ht16* mutants. (C) Single fruit weight, Yield per plant (3 fruits per plant), SSC of fruits, fruit length, fruit diameter, and carpopodium length of wt and *ht16* mutants’ fruit (final harvest period). (D) Dynamic data on fruit development of wt and *ht16* mutants. (E) Cell state and size in the longitudinal section of the fruits (16 DAA, near to the main vascular bundle). DAA, day after anthesis; L/D, ratio of fruit length to fruit diameter; two-way arrow in (A) point to carpopodium; The arrow in (D) point to the cell wall. Bars in (A) = 5 cm. Bars in (D) = 100 μm. *P < 0.05; ***P<0.001.

The dynamic development of fruits from 0 to 22 DAA was also tested. It turned out that the differences in fruit length between *ht16* and wild types was already evident in the early stage of fruit development, while the differences in fruit diameter only appeared at later developmental stages (Fig. 5D). The ratio of fruit length to fruit diameter (L/D) also showed significant differences in the middle and late stages of development (Fig. 5D). The decreased size of vertical cells in the *ht16* fruit is the main factor leading to the shortening of this *ht16* organ, while an increased number of transverse cells in the fruit is the main factor leading to the thickening of this organ (Fig. 5E).

To further elucidate why the *ht16* mutant fruit becomes shorter but bigger, we performed transcriptome analyses on 9 DAA fruits (*ht16* and wild types). The comparison of the gene transcripts in both genotypes showed that knocking out of *CsHT16* resulted in a significant decrease in the expression of *SWEET* genes, especially the *SWEET7* (Fig. 6B, Supplemental Fig. S6A), which gene product has been reported to mediate phloem unloading in companion cells which is required for proper fruit development (Li *et al*., 2021). Accompanied with a reduced expression of *CsHT16* and *CsSWEET7* a decrease of the glucose concentration in fruit flesh cells was observed, which is accompanied with altered transcripts of genes involved in phytohormone signaling (Fig. 6). E.g., there are some differentially expressed genes (DEGs) involved in either biosynthesis or responsive to auxin (Fig. 6C), representing a hormone requited for cell expansion. All DEGs involved in zeatin biosynthesis, a hormone affecting cell division, are down regulation (Fig. 6D). Similarly, all DEGs involved in brassinosteroid biosynthesis, a hormone affecting cell elongation, are also significantly down regulated (Fig. 6E and Supplemental Fig. S5A). With exception of the modified *CsHT16* gene, all other *CsHT* genes in fruits showed no significant difference in their respective transcript levels (Supplemental Fig. S5B).

**Fig. 6.**
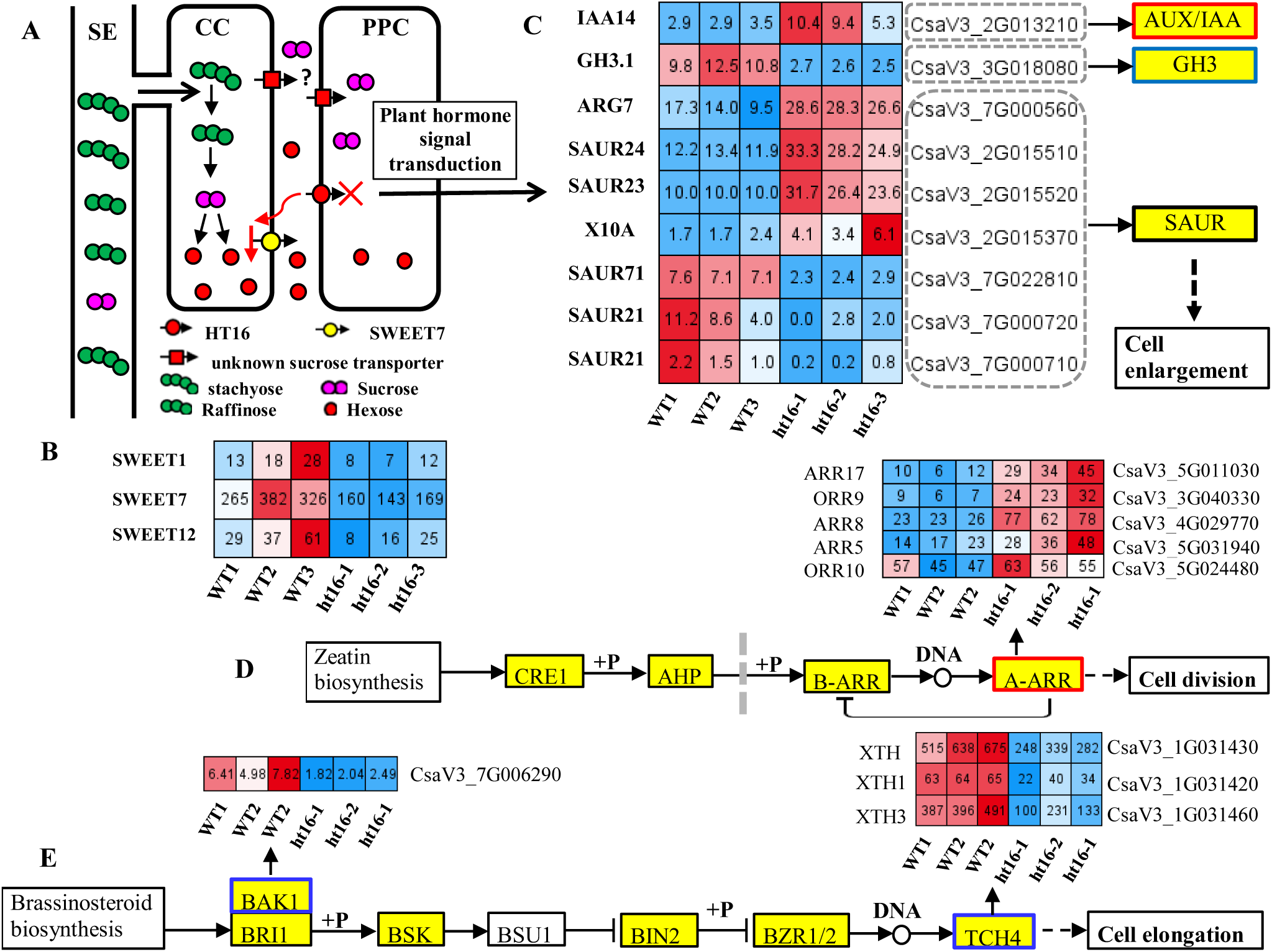
Knocking out of *CsHT16* causes changes in the plant hormone signaling pathway in cucumber fruits. (A) A working model of the impact of *CsHT16* knockout on sugar unloading in the phloem of cucumber fruits, *Cs*HT16 act as a hexose transporter located to plasma membrane of PPC, responsible for transporting hexose from apoplatic space to PPC cells. When the function of *Cs*HT16 was block, the expression of *CsSWEET7* was significantly down regulated. Reduced hexose in PPC causes changes in auxin signaling in fruits. (B) Significantly downregulation of SWEET genes expression in *CsHT16* knockout lines; (C) – (E) Significant upregulation of gene expression related to cell enlargement (D), cell division, and cell elongation (E) pathways; The heatmap with red indicating higher expression levels and blue indicating lower expression levels; The data on the heat map grid represents the FRKM value. Abbreviation: SE, sieve element; CC, companion cell; PPC, phloem parenchyma cell.

### Simultaneously downregulating the expression of *CsHT3/12/16* affects fruit development

Based on previous information (Cheng *et al*., 2015a, 2019) and the results above concerning the spatiotemporal expression analysis, heterologous functional expression and transport studies on bakeŕs yeast cells, subcellular localization, and immunolocalization analyses we propose that *CsHT3*, *CsHT12*, and *CsHT16* are the most important hexose transporters during cucumber fruit development.

To test the role of *Cs*HTs in cucumber fruits in more detail we generated both, a *CsHT3* overexpression (OE) construct, in which the gene is set under control of the CaMV35S promoter, and RNAi-*CsHT* construct directed against transcripts coding for *Cs*HT3, *Cs*HT12 and *Cs*HT16. We generated 10 independent OE-*CsHT3* lines (T1) and two independent RNAi-*CsHT* lines (T1) from which one RNAi line and two independent OE lines were selected for the subsequent generation. The *CsHT3* expression significantly increased in the OE-*CsHT3* lines with no effects on *CsHT12* and *CsHT16* transcripts (Fig. 7A). In the selected RNAi-*CsHT* line, the expression of *CsHT3*, *CsHT12,* and *CsHT16* were all significantly decreased (Fig. 7A).

**Fig. 7.**
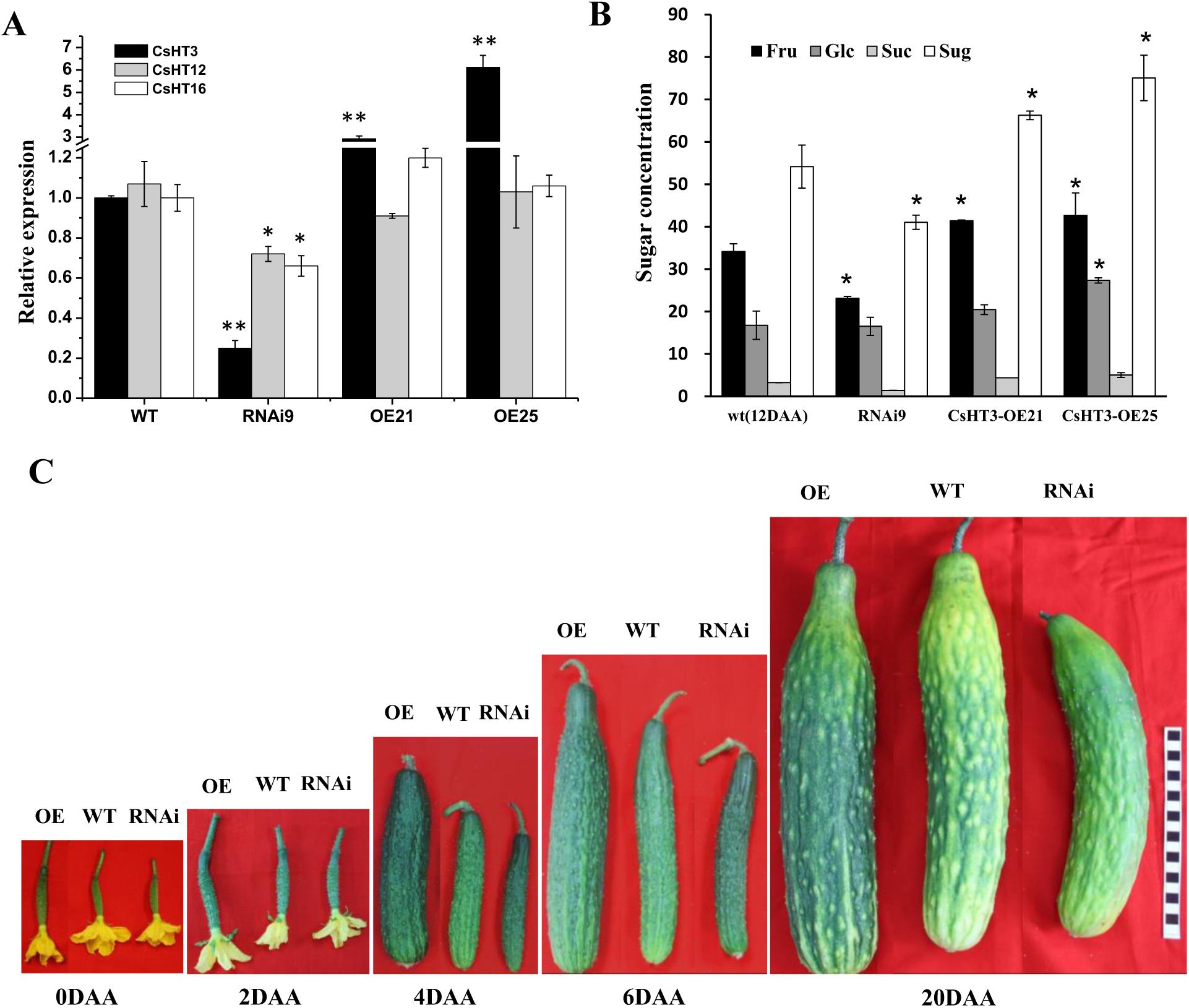
Biological function analysis of *CsHT* in cucumber fruit. (A) qRT-PCR detect the relative expression of *CsHT* in transgenic cucumber lines. (B) Sugar concentration analysis in *CsHT* transgenic fruits; (C) Phenotype analysis of transgenic cucumber fruits. OE, *CsHT3* over expression lines; RNAi, *CsHT* RNAi lines; Fru, fructose; Glc, glucose; Suc, sucrose; Sug, sugar; Bar in (C) = 1cm. The results represent at least 2 independent biological repetitions.

The fruit development of all transgenic plants was examined and it turned out that OE-*CsHT3* fruits grew faster and were larger than fruits of corresponding wild-types, while fruits from RNAi-*CsHT* mutants developed slower and were reduced in size (Fig. 7C, Supplemental Table S3). Accompanied with these morphological changes, fruits from mutants showed altered sugar levels at 9 DAA (corresponding to marketable mature stage). The levels of glucose, fructose, and sucrose appeared to be significantly increased in the OE-*CsHT3* fruits, while fructose contents were significantly decreased in fruits from RNAi-*CsHT* plants, when compared to wild types (Fig. 7B).

## Discussion

### Hexose transporters in cucumber

So far, three sugar transporter families are found all genomes from vascular plants, namely the Monosaccharide Transporters (MST), the Sucrose Transporters SUT/SUC) and the Sugar Will Eventually be Transported (SWEET) family (Pommerrenig *et al*., 2018). It has been shown that SUT/SUCs and SWEETs usually play roles in phloem loading (Chen *et al*., 2012; Wang *et al*., 2015), but their activities also impacting phloem unloading (Mizuno *et al*., 2016; Milne *et al*., 2017) and plant stress tolerance (Ma *et al*., 2018; Phukan *et al*., 2018).

MST proteins represent a large family with more than 50 members classified in 7 sub-families, including the HT/STP subfamily (Pommerrenig *et al*., 2018). *HTs/STPs* constitute a multigene family and the corresponding proteins are involved in sugar distribution and hexose transport in different tissues and different cell types (Büttner & Sauer, 2000; Slewinski, 2011). In Arabidopsis one of the seven subfamily members, namely the STPs, represent the best characterized hexose transporters. All of them locate to the plasma membrane and catalyze proton-driven hexoses import from the apoplastic space into the cell (Büttner, 2007).

In cucumber we identified in total 16 *HTs* genes that are homologous to the Arabidopsis *STP* genes (Supplemental Table S2). These cucumber genes are distributed on chromosomes 1 to 6; as no *HT* gene is present on chromosome 7 (Supplemental Fig. S2). Evolutionary tree analyses indicated that *CsHT1* and *CsHT11* are most closely related to *AtSTP9*, *AtST10*, and *AtSTP11* (Supplemental Fig. S1), and in-depth expression analyses revealed that these three Arabidopsis homologs showed, similar to *CsHT1* and *CsHT11*, pollen- or pollen tube-specific expression patterns (Schneidereit *et al*., 2003, 2005; Cheng *et al*., 2015b; Rottmann *et al*., 2016; Wen *et al*., 2020). The phylogenetic analysis demonstrated in addition that *CsHT12* and *CsHT16* are highly homologous with each other and closely related to *AtSTP1*. Moreover, *CsHT3*, *CsHT4,* and *CsHT15* are homologous to *AtSTP5*, *AtSTP14*, and *AtSTP7*, respectively (Supplemental Fig. S1).

Inspection of the *Arabidopsis thaliana* Genechip Database indicate that *AtSTP1, AtSTP5, AtSTP7, AtSTP12*, and *AtSTP14* are expressed in different stages of seed development (Büttner, 2010), implying that the highly homologous genes in cucumber (*HT3*, *HT4*, *HT12*, *HT15*, and *HT16*) may also impact fruit or seed development. However, our spatiotemporal expression analyses indicate that only *CsHT1* and *CsHT11* can be detected in the male flower tissue (Fig. 1), which represents an observation that is consistent to previous data (Cheng *et al*., 2015b). Among the *HT* in cucumber only *HT3*, *HT4*, *HT9*, *HT12*, *HT15* and *HT16* transcripts were detectable fruits, which is consistent with the information inferred from the phylogenetic tree *At*STPs have been described as hexose/H^+^ symporters which locate to the plasma membrane (Büttner, 2007) and our previous research showed that *Cs*HT1 and *Cs*HT3 also locate to the plasma membrane proteins and are able for glucose and galactose transport, while *Cs*HT4 is a galactose-specific plasma membrane protein (Cheng *et al*., 2015a). The transport experiments on *Cs*HT9, *Cs*HT12, and *Cs*HT16 revealed that all three carriers accept glucose, fructose, galactose and mannose as substrates, while *Cs*HT15 exhibited no transport activity for glucose or galactose, but activity when fructose or mannose are present as substrates (Fig. 2). This broad substrate spectrum of *Cs*HT9, *Cs*HT12, and *Cs*HT16 is in line with similar observations of the Arabidopsis homologs *At*STP1, *At*STP2, *At*STP4, and *At*STP6 (Sauer *et al.,* 1990; Truernit, 1996; Truernit *et al*., 1999; Scholz-Starke *et al*., 2003).

### Role of hexose transporters in post-phloem transport of cucumber fruit

Hu et al. (2011) have shown that phloem unloading in cucumber fruit follows an apoplastic route, indicating that sugar transporters are required for the subsequent uptake of sugars into the parenchymal cells from the apoplasmic space. It is known mature cucumber fruits mainly accumulate glucose and fructose (approximately 90% of the total soluble sugar) in the mesocarp and endocarp *(Handley et al., 1983; Hu et al.*, 2009) and this accumulation of these monosaccharides is consistent with the high activity of the cell-wall invertase during fruit development (Hu *et al*., 2011). Based on these findings we speculated that hexose transporters are the main sugar transporters involved in the post-phloem transport in cucumber fruits, which concurs with the molecular identification of two first hexose transporters in cucumber fruits *(Cheng et al.*, 2015a).

Here, our genome-wide base search for all the *HTs* in cucumber and the subsequent analysis of their expression patterns in fruits showed that apart from the two identified genes (Cheng *et al*., 2015a), four transcripts coding for further *HT*s are present in cucumber fruits (Fig. 1). From these four genes two the two newly identified, namely *HT12* and *HT16*, and one previously identified gene (*HT3*) exhibited the highest transcript levels in cucumber fruit (Fig. 1) (Cheng *et al*., 2019).

Our immunohistochemical localization studies showed that *Cs*HT3 and *Cs*HT16 were located in the parenchyma cells during fruit development (Fig. 4) which consists with the expression pattern of the gene encoding the cell-wall invertase *cwINV*, which has also been proven to locate to the SE/CC complex and in adjacent parenchyma cells of cucumber fruits (Hu *et al*., 2011). In sum, latter findings support our speculation that hexose transporters are involved in phloem unloading and post-phloem sugar transport in cucumber fruits. Immunohistochemical localization also showed that during ovary development, *Cs*HT3 was mainly located in CCs (Fig. 5). However, although *Cs*HT3 is described as a proton-driven sugar transporter (Cheng *et al*., 2015a) these localization des not rule out that this carrier may act as a sugar exporter. This is, because the proton-driven, phloem located sucrose transporter *Zm*SUT1, which is expressed in developing maize seeds, is also able for sugar export which is possible because of specific chemi-osmotic conditions in sink tissues (Carpaneto *et al*., 2005). Such transport of HT3 may even explain why this carrier is involved in development of the ovary phloem (Cheng *et al*., 2015a).

In general, apoplasmic unloading occurs in sinks which accumulate sugars at high concentrations (Lalonde *et al*., 2003); such as fruits from apples (Zhang *et al*., 2004), grape (Zhang *et al*., 2006) and tomato (Ruan & Patrick, 1995). Unexpectedly, sugar export in cucumber fruits (0–9 DAA) follows the apoplastic pathway although the total sugar levels are quite low (Hu *et al*., 2009, 2011). We speculate that this strategy is related to the translocation of stachyose and raffinose in cucumber plants and the rapid growth of the corresponding fruit. Cucumber, a typical stachyose and RFO-transporting plant (Hu *et al*., 2011) produces fruits which exhibit a remarkable fast growth rate (Ando & Grumet, 2010) and this fast growth requires high level of carbohydrates to support this process. In fact, the biomass of cucumber fruits increased more than 20-fold from 2 DAA to 4 DAA (Fig. 6c, Table 1) and the expression of *CsHT3*, *CsHT12*, and *CsHT16* increased sharply at 3 DAA of the fruit (Fig. 1D), supporting the proposed involvement of these carriers in post-phloem transport.

In line with this assumption we found that the reduced transcript levels of *CsHT3*, *CsHT12*, and *CsHT16* in our RNAi lines led to a decreased fruit enlargement and less sugar accumulation (Fig. 7) while on the other hand, *CsHT3* overexpression provokes increased fruit size and higher sugar accumulation during development of this organ (Fig. 6). These opposite results raised on over-expressors and RNAi mutants, respectively indicate that *Cs*HT3, *Cs*HT12, and *Cs*HT16 may play important roles in maintaining sink strength of cucumber fruits. This assumption receives independent support from the notion that cell-wall invertase, which hydrolyzes sucrose to hexose monomers, are required for an efficient phloem unloading in fruits by maintaining the sucrose gradient between the start of transport phloem (leaves) and the unloading sites (Patrick *et al*., 2001). Thus, in case of cucumber the hexoses present in the apoplasmic space will be rapidly imported into phloem parenchymal cells by the hexose transporters *Cs*HT3 and *Cs*HT16 (Fig. 4). That HT3 and HT16 are involved in sugar import, sub-sequent to sucrose export from the phloem, is further supported by the observation that both proteins shift their SC-CC specific localization in young fruits to a phloem parenchyma localization in developing fruits (Figs. 1 and 4), which are in the phase of rapid growth (Fig. 5). In line with this the knock down of *CsHT3*, *CsHT12* and *CsHT16* expression suppresses fruit growth (Fig. 7).

### The role of hexose transporters in maintaining cellular glucose homeostasis and sugar signaling

Glucose has multiple roles in plants as it serves e.g., as energy source, building block for anabolic processes, as osmotic regulator, and also as a potent signaling molecule that regulates gene expression controlling growth and development (Rolland e*t al.*, 2006; Cho & Yoo, 2011). We observed that impaired activity of *Cs*HT16, which according to our transport assays (Fig. 2) and localization studies (Figs. 1 and 4) transports glucose and fructose into fruit phloem parenchyma cells, induces changes of fruit morphology without affecting fruit biomass (Fig. 5). Through transcriptome sequencing analysis it was found that the expression of genes involved in the cell expansion pathway (auxin signaling related) and cell division pathway (cytokinin signaling related) were significantly increased in the fruit of the *ht16* mutant (Fig. 6). This may be due to the loss of *Cs*HT16 function, which leads to a decrease in glucose content in the parenchyma cells of the fruit phloem and flesh cells, causing changes in hormones that interact with glucose signals, leading to fruit thickening. In fact, in various studies a clear interaction between sugar levels and auxin- and cytokinin signaling has been confirmed (Rolland *et al*., 2006; Mishra *et al*., 2009).

We also found that the expression levels of multiple xylolucan endotrans glucose/hydroxylase protein (XTH) genes were significantly downregulated in the fruit of the *ht16* mutant (Fig. 6), and the expression of these genes has also been reported to be closely related to cell elongation. Taking Arabidopsis as are representative species it was show that the expression of *XTH18*, *XTH22*, and *XTH24* was suppressed by the SHORT-ROOT transcription factor to controls hypocotyl cell elongation (Dhar *et al*., 2022) and the activation of XTH17 and XTH24 is under control of LONGIFOLIA genes to promote polar cell elongation (Lee *et al*., 2018). Therefore, the reason for the shortened fruit of the *ht16* mutant may be due to a significant decrease in *XTH* gene expression. The XTHs are a family of enzymes that specifically use xyloglucan as substrate and is involved in cell wall extension process (Yokoyama *et al*., 2004; van Sandt *et al*., 2007). As glucose is the basic precursor for synthesizing xyloglucan synthesis (Kim *et al*., 2020) the mutation of *Cs*HT16 leads to a decrease in the content of glucose and other hexoses in cucumber fruit cells, which may affect the synthesis of xyloglucan and lead to a decrease in XTH gene expression (Fig. 6).

With the molecular identification of various *Cs*HT proteins critical the cucumber fruit development and impacting fruit properties, we provide insight into sugar homeostasis characteristics affecting the growth rate of a fast-developing fruit. These insights may even become instrumental for directed breeding approaches, in order to improve quality and yield of cucumber fruits.

## Materials and methods

### Plant materials

Cucumber (wild-type: *Cucumis sativus* L. cv. Xintaimici and transgenic lines) plants were grown under plastic house conditions from March to June or August to November in Wuhan (central China). Tissues were sampled for testing of gene transcription and sugar quantification. Different tissues (root, stem, leaf, petiole, female flower, male flower, and fruit) were collected from 2-month-old wild-type plants and used for spatial expression. Meanwhile, cucumber fruits at different developmental stages (stages 9–12; 0, 3, 6, and 9 DAA) were harvested for temporal expression and immunolocalization. All tissue and fruit samples were immediately frozen in liquid nitrogen and stored at −80 °C before use.

### RT-PCR and qRT-PCR

RNA extraction and reversion were based on the report of Cheng et al (2018). Supplemental Table S1 lists the primers for RT-PCR or qRT-PCR analyses. *TUA* was used as the control gene. SYBR green (TaKaRa) and ABI7500 system (Bio-Rad) were used to perform qRT-PCR. The mean expression level of relevant genes was calculated by 2^−ΔΔ^ ^Ct^ method (Livak & Schmittgen, 2001).

### Subcellular localization

To analyze the subcellular localization of CsHT9/12/16, we generated *CsHT*9-/12-/16-GFP fusion constructs by using the vector pH7LIC5.0-ccdB rc-N-eGFP with the primers listed in Supplemental Table S1. The constructs were used to inject *N. benthamiana* with p19, as described by Cheng et al (2018). The mCherry-labeled construct PM-rk (CD3-1007) was used to co-express with GFP as the PM. Transient expression of the fusion protein CsHT16-GFP in cucumber protoplast was conducted as described by Cheng et al (2015a). Fluorescence was imaged using the FV1200 Olympus confocal laser scanning microscope.

### Heterologous expression analysis of *Cs*HTs in yeast

The ORFs with appropriate restriction sites of *CsHT9, CsHT12, CsHT15*, and *CsHT16* were constructed into the yeast expression vector pDR196 with the primers listed in Supplemental Table S1 and then transfected to the hexose transporter-deficient yeast (*S. cerevisiae*) strain EBY.VW4000 (Wieczorke *et al*., 1999) according to the method of Morita & Takegawa (2004). The drop test for yeast growth and metabolic inhibitor treatment were performed according to the work of Cheng *et al*. (2015a).

For the time course of yeast growth, the transformed cells were pre-grown in liquid SD-URA medium supplemented with 2% (w/v) maltose as the sole carbon source to an OD_600_ of approximately 0.8. Cells were washed with ddH_2_O and suspended in the medium with glucose as sole carbon source to an OD_600_ of 0.1. After growing at 30 °C, the OD_600_ data were collected at 4, 6, 8, 10, 12, and 14 h. For the pH dependence of yeast growth, the transformed cells were also pre-grown in medium with maltose as sole carbon source to an OD_600_ of approximately 0.8. After washing with ddH_2_O, the cells were suspended in the glucose medium with different pH values (2, 3, 4, 5, 6, 7, and 8). After growing at 30 °C, the OD_600_ data were collected at 8 h.

### Immunolocalization of *Cs*HT3 and *Cs*HT16 in cucumber fruit

The CsHT3-antibody has been prepared previously, as shown in our previous paper (Cheng *et al*., 2015a). For CsHT16, two specific peptide fragments (MPAVAAIVPGDTKKEYPC and CTHWYWSRFVTDNNFQIG) were selected to synthesize polypeptides and used for immunization of two rabbits (specific pathogen-free) by Beijing B&M Biotech.

For immunohistochemical analysis, cucumber fruits at different developmental stages were sliced into transverse paraffin sections (14 µm) and used as previously described (Cheng *et al*., 2015a, b). The specimens were viewed with a Nikon eclipse 80i microscope.

### Construction of expression vector and generation of transgenic cucumber plants

A specific sgRNA (GACACGTAAGTTCGGTCGT) of CsHT16 was selected by using Geneious _Prime and the off-target analysis was carried out on a website (http://www.rgenome.net/cas-offinder/). The sgRNA of CsHT16 was constructed into the PKSE402 vector which was constructed by inserting 35S:GFP:Terminator into the EcoRI site of PKSE401(Xing *et al*., 2014; Hu *et al*., 2017). To construct the *CsHT3* overexpression vector, we isolated the coding region of *CsHT3* from Xintaimici cucumber fruit through PCR. The DNA was digested with XbaI/SmaI and cloned into the PBI121 vector under the control of the CaMV *35S* promoter. For RNAi construction, a 111 bp fragment from the coding region of *CsHT3* cDNA (bases 74– 184) was amplified by PCR and introduced in sense and antisense orientations into the RNAi vector pFGC1008 under the control of the CaMV 35S promoter to generate RNAi-*CsHT* construct. Then, the vectors were transformed into *Agrobacterium tumefaciens* LBA4404. The primers used are listed in Supplemental Table S1. The *A. tumefaciens* with CsHT3 overexpression or RNAi-*CsHT* vector was transformed into the cucumber pure line Xintaimici by using fresh expanding cotyledon disk transformation, as previously described (Cheng *et al*., 2015b; Hu *et al*., 2017). The regenerated plants were screened by RT-PCR analysis.

After several generations of self-pollination, we obtained homozygous trans-free ht16 gene-edited lines. Phenotype identifications were confirmed in both 2021 and 2022 in plastic house, and adequate plants were used for fruit morphology analysis.

### Quantification of soluble sugars

The sugar content in the transgenic cucumber fruit was determined by quantitative GC-MS (gas chromatography–mass spectrometry), as described by Cheng *et al*. (2018). The 12 DAA fruits from different transgenic cucumber and wild-type plants were collected, and the mesocarp tissues were sampled for sugar determination.

### RNA-seq analysis

Fruit (9 DAA) flesh samples from WT and ht16 were collected and stored at 80 ℃ after freezing with liquid nitrogen. Total RNAs of these samples (3 replications for each treatment) were isolated using TRIzol reagent (Invitrogen), respectively. High-quality RNA samples were used to construct sequencing libraries with NEBNext Ultra RNA Library Prep Kit for Illumina (NEB, USA), and the libraries were sequenced using Illumina HiSeq X ten/NovaSeq 6000 system (2 × 150 bp read length) at Majorbio Biological Technology (China) (https://www.majorbio.com/). Each sample yielded more than 6 gigabytes of data and clean reads were mapped to reference cucumber genome Chinese Long v3.0 via TopHat2 (http;//tophat.cbcb.umd.edu/).

Gene expression levels were estimated by fragments per kilobase of transcript per million fragments mapped (FPKM). Genes with more than a twofold change at the expression level and P value < 0.05 were set as the threshold for significantly differential expression (DEG).

### Bioinformatics analysis

Primers were designed using the Primer Premier 5.0. Sequence homology search in GenBank was performed with the BLAST program (http://www.ncbi.nlm.nih.gov/BLAST/; Madden et al. [1996]).

Phylogenetic analysis was conducted using MEGA version 5.2 (Kumar et al., 2004), adopting Poisson correction distance, and it was presented using traditional rectangular TreeView. Support for the tree was assessed using the bootstrap method with 1,000 replicates.

### Statistical analysis

Student’s *t*-tests were performed using the algorithm embedded in Microsoft Excel, and significance was evaluated at the 5% level (*P* < 0.05) for all comparisons. For each treatment, the standard error of the mean (SE) was calculated based on at least three biological replicates.

## Acknowledgments

We thank Dr. Xiaolei Sui (China agricultural university) for the gifts of CsHT3-antibody and yeast strain EBY.VW4000. This work was supported by the National Natural Science Foundation of China (31972435 and 31601774) to Jintao Cheng. Natural Science Foundation of Hubei Province (2019CFA017), and the Fundamental Research Funds for the Central Universities (2662023YLPY008). Work in the lab from E. Neuhaus was supported by the Forschungsinitiative Rheinland-Pfalz. BioComp.

## Author contributions

JC and ZB designed the experiments. JC and SW performed most of the experiments. KL, YZ, and MZ planted the transgenic plants and counting phenotypes. JC and SW analyzed the data. JC wrote the article. SW, ZB, HEN and YZ read and revised the article.

## Conflict of interest

The authors declare no conflict of interest.

## Data availability statement

All relevant data can be found within the manuscript and its supporting materials.

## Supporting information

Additional Supporting Information may be found in the online version of this article.

## Supplemental Data List

**Supplemental Fig. S1** Phylogenetic tree analysis of *Cs*HTs and AtSTPs. Database accession numbers of the sequences used are as follows: *Cucumis sativus* CsHT1(HQ202746), *Cs*HT2(KP113693), *Cs*HT3(KP113694), *Cs*HT4(KP113695), *Cs*HT5 (XP_004141211.1), *Cs*HT6 (XP_004134451.1), *Cs*HT7 (XP_004134452.1), *Cs*HT8 (XP_011652758.1), *Cs*HT9 (XP_011657575.1), *Cs*HT10 (XP_004134450.2), *Cs*HT11 (XP_011654900.1), *Cs*HT12 (XP_004146580.1), *Cs*HT13(XP_004146944.1),_____*Cs*HT14 (XP_004143993.1),*Cs*HT15 (XP_004146734.1), *Cs*HT16 (XP_011658192.1); *A. thaliana At*STP1 (NM_100998), *At*STP2 (NM_100608), *At*STP3 (AJ002399), *At*STP4 (NM_112883), *At*STP5 (AJ344335), *At*STP6 (AJ344337), *At*STP7 (AJ344331), *At*STP8 (AJ344332), *At*STP9 (NM_103915), *At*STP10 (NM_112884), *At*STP11 (AJ001664), *At*STP12 (AJ344333), *At*STP13 (AJ344338), and *At*STP14 (AJ344334).

**Supplemental Fig. S2** Distribution of *CsHTs* in cucumber chromosome.

**Supplemental Fig. S3** Subcellular localization of *Cs*HT9 and *Cs*HT12 in tobacco protoplast. The laser scanning confocal microscopy images show fluorescence and merged images. The bright field is also presented. GFP, green fluorescence protein; RFP, red fluorescence protein; Bar = 20 µm.

**Supplemental Fig. S4** Pre-immune serum was used as the control in Figures 4(A) and (C). Bar =100 µm.

**Supplemental Fig. S5** qRT-PCR detected the expression of some DEGs that were shown in Figure 6 (A) and FPKM of *CsHTs* in fruits. FPKM: Fragments Per Kilobase of exon model per Million mapped fragments.

**Table S1** Primers used in the paper (Lowercases indicate the restriction enzyme sites or recombination connector sequence).

**Table S2** Sequences information of *CsHTs*

**Table S3** Transgenic cucumber fruit size statistics

